# Structural Effects of Low Social Status and Obesogenic Diet on Social and Emotional Neurocircuits in Female Macaques: A Longitudinal Study from Infancy to Adulthood

**DOI:** 10.1101/2025.07.30.667163

**Authors:** Z. Kovacs-Balint, A. Gopakumar, M. Kyle, J.R. Godfrey, K. Bailey, T. Jonesteller, A.C. Gray, K. Shabbir, A. Wang, J. Acevedo, R. Vlasova, M. Styner, J. Raper, J. Bachevalier, M.C. Alvarado, K Ethun, ME Wilson, M.M. Sanchez

## Abstract

A substantial body of literature has demonstrated a consistent link between psychosocial stress and obesity in children, particularly in those from low socioeconomic backgrounds. Despite evidence indicating a complex interplay between stress, diet and obesity, there is a limited understanding of the specific versus potentially synergistic effects of obesity and stress on brain structural and functional development. This study investigates the developmental and long-term brain structural alterations resulting from exposure to chronic social stress due to low (subordinate -SUB-) social status and postnatal obesogenic diets. Forty-one female rhesus macaques (Dominants -DOM-, n=21; Subordinates -SUB-, n=20) were assigned to either only low-calorie diet (LCD) or to both high-calorie diet (HCD) and LCD (Choice diet) from birth through the juvenile period. After menarche, all subjects were maintained on a LCD-only diet through adulthood. Twenty-seven animals (DOM: n=13, SUB: n=14) were studied again in adulthood to investigate the long-term effects of early diet and social rank on brain structure. Cumulative Kcal consumption was measured from birth through 16 months and body weights were measured at all time points.

Overall, the findings show specific effects of obesogenic diet and psychosocial stress on cortical and corticolimbic brain regions. Animals with access to the obesogenic diet had larger overall brain size (measured as intracranial volume -ICV-) and larger overall volumes of prefrontal cortex, insula, superior temporal sulcus (defined as temporo-parieto-occipital area rostral and caudal regions (TPOr and TPOc)) than those in the low-calorie diet. Most of these regional diet effects, except for the insula, were driven by general effects of the diet on brain size. The diet effects were lost when adding the adult data to the longitudinal analysis, suggesting transient effects of obesogenic diets while the animals were consuming it, but not long-term, persistent effects. These findings highlight the potential of brain rescue mechanisms that could offset lasting developmental effects of early-life obesogenic diet consumption. With respect to social rank, SUB exhibited larger volumes in brain regions related to social cognition and emotional processing than DOM animals. When the adult data was added to the longitudinal analysis, the effects of social rank were prominent in the hippocampus, superior temporal sulcus, temporo-parieto-occipital rostral region, and the temporal auditory cortices after ICV data correction, suggesting long-term, persistent and cumulative effects of these social experiences, in contrast to the transient diet effects.

## 1. Introduction

Early life adversity/stress (ELA/ELS) is a risk factor for physical and mental illness throughout the lifespan (Green et al., 2010; McLaughlin et al., 2012) and has become a critical target of public health initiatives. ELA includes a broad range of adverse experiences such as childhood maltreatment, exposure to domestic violence, bullying, trauma exposure and bullying (Wade, Wright, & Finegold, 2022). Children and adolescents exposed to ELA have a heightened risk of developing psychiatric and behavioral disorders, including depression, conduct disorder, substance use, psychosis, self-harm behaviors, and suicide (Danese, McLaughlin, Samara, & Stover, 2020; El-Khodary & Samara, 2020; Schaefer et al., 2018). Consistent with alterations in emotional and stress regulation, structural and functional alterations have been reported in prefrontal cortex (PFC)-amygdala-hippocampus circuits of youth with ELA living in environments plagued with constant threat and adversity (McLaughlin et al, 2019); these include increased amygdala (AMY) activity to negative stimuli (McCrory et al., 2013; McCrory et al., 2011; McLaughlin, Peverill, Gold, Alves, & Sheridan, 2015; White et al., 2019), reduced volumes in medial prefrontal cortex (mPFC) and hippocampus (van Harmelen, van Tol et al, 2010; Edmiston, Wang et al, 2011). These brain alterations have been identified as risk factors for the development of depression, anxiety, and general psychopathology (Hein & Monk, 2017), further underscoring the urgency to understand how ELA/ELS negatively impacts neurobehavioral development.

In addition to long-lasting mental illness and emotional turmoil, ELA and chronic stress exposure during childhood can lead to negative physical health outcomes such as cardiometabolic disease, obesity, and chronic inflammatory processes (Drewnowski, 2009; Evans & English, 2002; Hemmingsson, 2018; Lipowicz, KozieŁ, Hulanicka, & Kowalisko, 2007; Ogden et al., 2018)-later in life if it exceeds a child’s coping abilities (Audage & Middlebrooks, 2008). These health disparities are frequently observed for environmental stressors embedded in conditions of low socioeconomic status -SES- (Audage & Middlebrooks, 2008; Wyman et al., 1992). Although the effects of SES on a child’s health is dictated heavily by the resources available (Winkleby et al., 1992), exposure to chronic stressors are critical contributors (Evans & English, 2002), including food insecurity, psychosocial stress and trauma, exposure to violence, and unstable home environment (Myers, 2020; Winkleby et al., 1992).

Low SES, psychosocial and ELA have also consistently been linked with childhood obesity. The prevalence of obesity among US children/youth (aged 2-19 years) is low in higher income groups compared with other groups (Ogden et al., 2018) and some studies have found a significant inverse association between family SES and obesity (Murasko, 2013). Low SES families experience lower incomes that limit access to high-nutrient foods and live in “food deserts” and neighborhoods that increase the likelihood of consumption of obesogenic diets (Case et al., 2002; Drewnowski & Darmon, 2005). In addition to limited access to resources low SES environments generate chronic exposure to stressors; and chronic stress has been shown to promote over-eating of highly caloric diets in humans (Adam & Epel, 2007; Greeno & Wing, 1994) and animal models (Foster, Solomon, Huhman, & Bartness, 2006; Godfrey et al., 2018; Hagan, Chandler, Wauford, Rybak, & Oswald, 2003; Solomon, Foster, Bartness, & Huhman, 2007), leading to increased risk for comorbid obesity (Evans, Fuller-Rowell, & Doan, 2012). Notably, females are more vulnerable to emotional and stress-induced over-eating, increased consumption of highly caloric (Western) diets, and obesity (Arce, Michopoulos, Shepard, Ha, & Wilson, 2010; Suglia, Duarte, Chambers, & Boynton-Jarrett, 2012).

Chronic stress can result in metabolic syndrome (including obesity) through elevations of stress hormones, inflammation, and glucocorticoid resistance (Macfarlane, Forbes, & Walker, 2008; Rosmond, 2005). Then, an increase in adipose tissue can contribute to chronic inflammation by way of enlargement of adipocytes and subsequent decreased blood supply, or hypoxia, to these cells, stimulating the release of pro-inflammatory adipokines (Ellulu, Patimah, Khaza’ai, Rahmat, & Abed, 2017). Increased peripheral inflammation can activate microglia in the brain, prompting release of local inflammatory cytokines which results in neuroinflammation (Shields, Haque, & Banik, 2020). During development, over-activation of microglia can cause alterations in synaptic pruning and impact myelination and brain white matter (Mottahedin et al., 2017), affecting brain structure and function. For both humans and animal models, chronic stress shifts eating preferences to high-calorie foods (Hill, Moss, Sykes-Muskett, Conner, & O’Connor, 2018), and, in the case of children, early exposure to obesogenic diets can affect dietary preferences and health outcomes as adults (Teegarden, Scott, & Bale, 2009). Altogether, this evidence provides potential biological and neural mechanisms by which chronic stress early in life and highly caloric, obesogenic diets, can work together to derail brain development and increase risk for neuropsychiatric disorders.

While there is evidence regarding the complex interplay between chronic stress, eating habits, and obesity, there is limited understanding of the specific versus potentially synergistic effects of chronic stress and obesogenic diet on developmental alterations in brain structure and function. The goal of this study was to examine the developmental and long-term, cumulative, impact of social subordination stress and early life exposure to an obesogenic diet on structural development of brain regions involved in emotional and stress regulation, sensory processing, and homeostasis in a translational nonhuman primate (NHP) model. NHPs, including rhesus macaques (*Macaca mulatta*), present a unique opportunity to study the specific and synergistic developmental and long-term effects of exposure to chronic psychosocial stress and highly caloric, obesogenic, diets on brain development with high experimental control. Leveraging the macaque model allows to circumvent challenges frequently encountered in human studies, such as limited control over environmental variables and diets, as well as confounding factors (e.g. drug abuse, education, healthcare access). Furthermore, the social subordination (SUB) model in macaque hierarchies and the use of automated feeders that allow to control animals’ access to different diets and measure Kcal intake while living in social groups, are well validated approaches to investigating the effects of chronic psychosocial stress (Wilson et al., 2008). Moreover, the rhesus monkey brain exhibits high similarities in neural circuity and developmental trajectories to the human brain, also longer relative to less sentient mammals, underscoring the translational potential of this model (Kovacs-Balint et al., 2021). Specifically, macaques and humans share striking similarities in brain structures related to social functioning including cortical regions such as the superior temporal sulcus (STS), prefrontal cortex (PFC), and hippocampus which support higher cognition (Isoda, Noritake, & Ninomiya, 2018). Macaques also share complex social behaviors with humans and have complex social structure and dominance hierarchies (Bernstein, 1976), making them a valuable model for analyzing brain structural MRI data across development.

Rhesus monkeys live in female societies with matrilineal and matriarchal social structures where infants assume their mothers’ social status and males have a separate social hierarchy (Bernstein, 1976). The hierarchical dominance hierarchies are maintained through aggression, where low status females receive more aggression and harassment from dominant animals, less affiliative behavior, and exhibit higher stress and anxiety levels than dominants (Carol A. Shively & Day, 2015; Wilson, 2016). Due to this dominance hierarchy, the incidence of fighting diminishes, as subordinates typically exhibit submissive behaviors -particularly withdrawal- and abstain from retaliatory physical aggression, thereby acknowledging their place in the social hierarchy (Bernstein & Gordon, 1974; Bernstein, Gordon, & Rose, 1974). This results in hypothalamic-pituitary-adrenocortical (HPA) axis over-activation and chronic-stress-related phenotypes in subordinates -e.g. emotional dysregulation and higher expression of genes related to inflammation- (Michopoulos, Higgins, Toufexis, & Wilson, 2012; Michopoulos, Reding, Wilson, & Toufexis, 2012; C. A. Shively et al., 2005; Wilson, 2016). The effects of social subordination are present early in development, with low-ranking animals starting to receive aggression during infancy, although not before weaning (Spencer-Booth, 1968). Thus, by the time subordinates are juveniles, they already show elevated levels of the stress hormone cortisol (Howell et al., 2014). Additionally, immunological and hormonal signals passed in breastmilk from subordinate mothers could also contribute to early onset of alterations in structural brain development in the infants -evidenced by measures of white matter integrity- (Casabiell et al., 1997; Howell et al., 2013). Given that social status remains relatively constant for females throughout their life, in contrast to males who are often placed in different social groups and encounter changes in social rank, only females subjects were utilized in this study (Wilson, 2016).

Earlier studies demonstrated that subordinate female macaques tend to consume more highly caloric foods (Arce et al., 2010). These behavioral patterns predispose subordinates to increased adiposity and obesity. Thus, social subordination not only elicits chronic stress that affects the brain, but also predisposes the animal to potential obesity-related complications by increasing the motivation to eat palatable, calorically dense diets (Arce et al., 2010). Infant rhesus monkeys depend on breast milk as their primary source of nutrition until approximately 6 weeks of age, when dams begin to introduce solid foods into their infant’s diet, in parallel to increased infant exploration, self-sufficiency and decreased contact with the mother (Hinde & Spencer-Booth, 1967). As the weaning process progresses between 3 to 6 months of age, they transition from a breast milk-based diet to a solid food-based diet, marking the reduction of breastfeeding (Reitsema, Partrick, & Muir, 2016; Stroud et al., 2006). The utilization of the rhesus macaque model of social subordination, in conjunction with experimental control of diet conditions, allows for a deeper understanding of how neurodevelopmental alterations happen, which can inform clinical interventions to mitigate the negative effects of chronic psychosocial stress and obesogenic diets.

Complex social behaviors are supported by a distributed neural network involved in the processing of social information, salience processing, somatosensory integration, socioemotional regulation, reward/motivation, homeostatic control and decision-making. In addition to the role of the hippocampus, cortical regions in this network include the prefrontal cortex (PFC), superior temporal sulcus -STS- (Bickart, Wright, Dautoff, Dickerson, & Barrett, 2011; Lewis, Rezaie, Brown, Roberts, & Dunbar, 2011; Noonan et al., 2014; Powell, Lewis, Roberts, García-Fiñana, & Dunbar, 2012; Sallet et al., 2011) and the insular cortex -INS- (Augustine, 1996; Lighthall et al., 2012; Singer, Critchley, & Preuschoff, 2009; Uddin, Nomi, Hébert-Seropian, Ghaziri, & Boucher, 2017). The PFC guides complex cognitive and emotional behaviors linking sensory inputs to motor outputs (Barbas & Zikopoulos, 2007). The medial PFC (mPFC) and the dorsolateral PFC (dlPFC) are involved in executive function and attention. Chronic stress during the juvenile stage of development can lead to reduced gray matter (GM) volume in key regions of the PFC, including the dlPFC, anterior cingulate cortex (ACC), and mPFC, and structural alterations in these regions have been linked to cognitive rigidity, increased emotional reactivity, and impaired socioemotional function (Arnsten, 2009). Specifically, chronic stress can lead to reduced dendritic arborization and branching of mPFC neurons that could contribute to the structural alterations in GM (McEwen & Morrison, 2013b). The impacts of stress on the PFC could be long-term, as evidenced by the reduced GM density in the PFC of adult male macaques who experienced social subordination since juveniles in comparison to dominant animals (Sallet et al., 2011). The superior temporal sulcus (STS), a key cortical area of the primate “social brain” (Boddaert et al., 2004; Ohnishi et al., 2000), responsive to facial expressions, movement, and eye-gaze, and develops rapidly within the first 18 months of life in humans (Allison, Puce, & McCarthy, 2000). Increased fMRI activation of the STS is detected during speech and language processing, and critical involvement in audiovisual processing of social information -including the perception of faces and human motion- and theory of mind (Deen, Koldewyn, Kanwisher, & Saxe, 2015; Raschle et al., 2014). Within the STS, the temporo-parieto-occipital (TPO) region is particularly interesting for social behaviors, because it contains polysensory neurons that show strong responses to combinations of visual, auditory, and/or somatosensory stimuli (Baylis, Rolls, & Leonard, 1987; Bruce, Desimone, & Gross, 1981). The TPO is connected with unimodal association areas involved in these three sensory modalities (Barnes & Pandya, 1992), suggesting that the TPO is important for high-level integration of sensory information, including perception of biological motion -“motion patterns characteristic of living organisms in locomotion”- (Johansson, 1973). Thus, the TPO plays an important role in processing and integration of complex social information in humans and macaques. In NHP studies, a positive correlation has been reported between social rank and GM volume in the mid-STS, inferior temporal gyrus, and temporal poles (Sallet et al., 2011; Von Der Heide, Vyas, & Olson, 2014). Furthermore, studies have shown that the macaque STS contains a distributed network of areas that are activated by the processing of gaze cues (Materna, Dicke, & Thier, 2008) with SUB engaging in more rapid gaze following than DOM animals (Shepherd, Deaner, & Platt, 2006). While it is evident that social rank influences STS activity, the neural mechanisms underlying this relationship are still being uncovered. Finally, the insular cortex (or insula -INS-) is a critical hub connected with multiple sensory, limbic and association regions that support its involvement in a range of physiological and cognitive functions essential for primates social interactions- (Augustine, 1996; Lighthall et al., 2012; Singer et al., 2009; Uddin et al., 2017). These functions include visceral sensing, limbic and somatosensory integration, motor activities, salience processing, interoception, socioemotional functions and decision-making (Evrard, 2019); Nieuwenhuys, 2012; Rachidi et al, 2021). The INS is composed of two distinct subregions, with their unique set of functions: posterior INS is primarily responsible for interoceptive, visceral, and physiological functions, while the anterior INS is more involved in socio-emotional functions and introspection of feelings (Chiarello et al., 2013). According to a prominent model of INS function, information flows from the posterior to the anterior pole, where the posterior INS plays a key role in the detection and integration of interoceptive signals related to bodily sensations, which are then integrated with emotional information in the middle INS and transferred to the anterior INS for self-awareness related to the sensory experience (Craig, 2009). The relationship between INS structural alterations and stress, however, is paradoxical. While the INS exhibits heightened activation across a range of anxiety and stress disorders, chronic stress has largely been associated with decreased INS volumes (Ansell, Rando, Tuit, Guarnaccia, & Sinha, 2012). Studies focusing on individuals with PTSD arising from ELA have revealed notable reductions in INS volume, with cumulative adversity displaying a correlation with INS GM volume loss (Ansell et al., 2012), and a meta-analysis showed INS hyperactivation in individuals with PTSD and/or social anxiety disorder (SAD) (Etkin & Wager, 2007); microglial overactivation, coupled with INS hyperactivity, may increase the INS susceptibility to excessive synaptic pruning and volume reduction (Eltokhi, Janmaat, Genedi, Haarman, & Sommer, 2020).

Interestingly, obesity and high caloric intake has been linked with some structural brain alterations that resemble those of chronic stress, including smaller insula and PFC volumes in adolescents (Gómez-Apo, Mondragón-Maya, Ferrari-Díaz, & Silva-Pereyra, 2021). Furthermore, studies have found negative correlations between body mass index (BMI) and temporal lobe volumes, where obese individuals had significant reduction in GM in the right temporal gyrus and left temporal pole (García-García et al., 2019; Opel et al., 2021). Associations between metabolic syndrome and decreased GM in the right superior temporal gyrus have been made as well (Kotkowski et al., 2019).

The goal of this study was to investigate the short- and long-term specific, and potentially synergistic effects of social subordination stress and an obesogenic diet on the structural development of brain regions responsible for stress and emotional regulation, socioemotional processing, reward, interoception, and homeostatic control. Specifically, we focused on PFC, hippocampus, temporal lobes (visual, auditory), STS, and the INS. To achieve this goal, we used a translational rhesus macaque model of social SUB compared to dominant (DOM) animals with subgroups assigned to either a low-calorie-only diet (LCD) or a “Choice” diet – consisting of access to both LCD and a highly caloric diet enriched in fat, and sugars (HCD). After the subjects reached puberty, they were placed on an LCD-only diet until adulthood to investigate the long-term, cumulative, effects of social rank in adults and whether the effects of the postnatal obesogenic diet persisted in the adult brain or whether the effects were transitory and rescued/normalized by the LCD diet.

## 2. Methods

### 2.1 Subjects, Housing, and Experimental Design

Brain development of 26 female rhesus macaques (*Macaca mulatta*) with different postnatal social status and diets from infancy through adulthood (2 weeks to ~7 years of age) were studied using magnetic resonance imaging (MRI). The animals were living in complex social environments comprised of 2-3 adult males, 30-60 adult females, and their offspring at the Emory National Primate Research Center (ENPRC) Field Station. Subjects were selected from different social dominance ranks (top third: Dominants -DOM, n=12-; bottom third: Subordinates -SUB, n=14-). Subjects were assigned to two different postnatal diets (low calorie diet -LCD-, or a “Choice” diet consisting of a high calorie diet -HCD- and LCD); and after menarche, all animals were assigned to the LCD-only diet condition. See breakdown of groups by social rank and diets in **Table 1**, and **Figure 1** for timeline, experimental design and measures collected longitudinally. Dams with a known history of neglect or physical abuse towards infants were excluded from the study. Infants, with a birth weight <450g, were also excluded from the study to avoid confounding effects of prematurity/low-birth weight on brain development (Aarnoudse-Moens, Weisglas-Kuperus, van Goudoever, & Oosterlaan, 2009).

**Figure 1.**
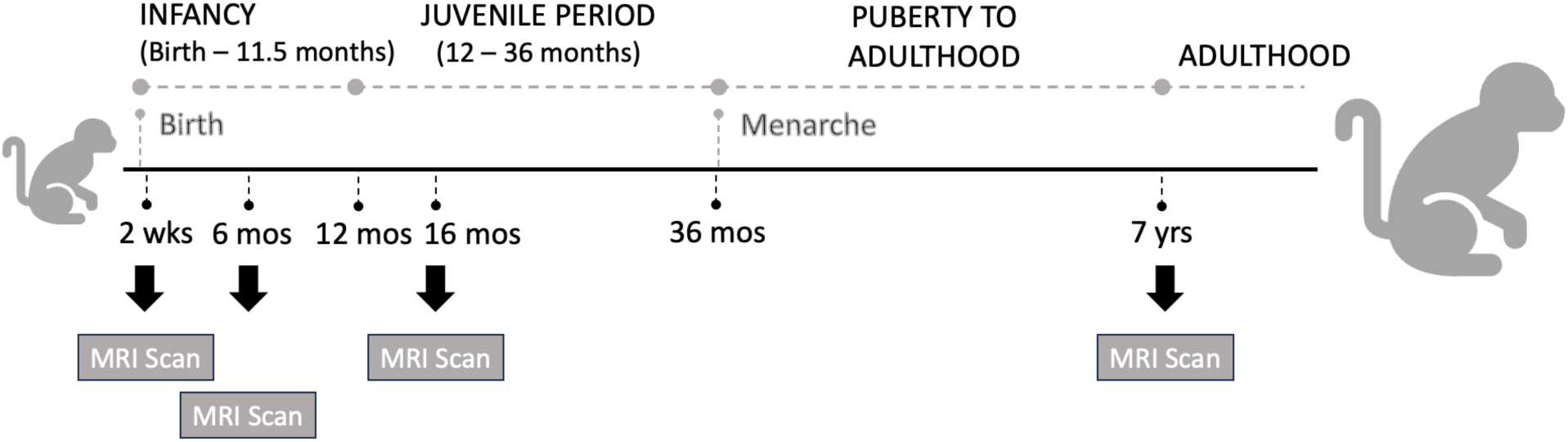
Timeline of the study. Twenty-six infant rhesus macaques were assigned after birth and were followed until adulthood. T1w and T2w brain MRI scans were collected longitudinally during infancy (at 2 weeks and 6 months), the juvenile period (at 16 months of life), and during adulthood (around 7 years of age).

**Table 1.** Breakdown of groups by social rank (dominant -DOM-, subordinate -SUB-) and diet (high+low-calorie diet -Choice-, only low-calorie diet -LCD-).

|  |  | **RANK** |  |  |
| --- | --- | --- | --- | --- |
|  |  | Dominant | Subordinate | Total |
| **DIET** | Choice | 5 | 8 | 13 |
|  | LCD | 7 | 6 | 13 |
| Total |  | 12 | 14 | 26 |

All procedures were approved by the Emory University Institutional Animal Care and Use Committee (IACUC), in accordance with the Animal Welfare Act and the U.S. Department of Health and Human Services’ “Guide for the Care and Use of Laboratory Animals.” Furthermore, the ENPRC is fully accredited by AAALAC International.

### 2.2 Diet Conditions

All pregnant females were fed a chow-based, LCD-only, healthy diet throughout pregnancy to avoid confounding effects of different prenatal dietary environments on our brain measures (Janthakhin, Rincel, Costa, Darnaudéry, & Ferreira, 2017; Sullivan et al., 2010). Mother-infant pairs were randomly assigned at birth to diet groups: LCD-only condition (n=7 DOM, 6 SUB) or to a Choice condition with access to both the LCD and HCD diet (n=5 DOM, 8 SUB) (Table 1). Previous research has established that, in comparison to a HCD-only diet, the Choice condition is a superior translational model for the “Western” human diet, allowing for stress-induced, emotional feeding in animal models and is shown to promote increased consumption of calorically dense foods (Michopoulos, Toufexis, & Wilson, 2012).

Subcutaneous radio-frequency identification (RFID) chips were implanted in the subjects’ wrists, allowing subjects’ access to and activation of automated feeders to deliver pellets of one of both of the two experimental diets and accurately record the Kcal consumption of each diet (Wilson et al., 2008).

These chips were programmed to enable each subject to access either only LCD feeders -LCD condition- or both LCD and HCD feeders -Choice condition-. This system has been validated at the ENPRC demonstrating that subjects almost always consume a pellet of food once they have taken it from the feeder (Wilson et al., 2008).

### 2.3 Structural MRI (sMRI) data

#### 2.3.1 sMRI Data Acquisition

T1- and T2-weighted sMRI scans were collected at 2 weeks, 6 months, 16 months, and during adulthood (7.01 ±0.1yrs of age) (Fig.1), using a 3T Siemens Magnetom TRIO system (Siemens Medical Solutions, Malvern, PA, USA) and an 8-channel phase array knee coil, following published sequences and protocols by our group for macaque brain (Godfrey et al., 2023; Kovacs-Balint et al., 2021; Kovacs-Balint et al., 2023; Reding, Styner, Wilson, Toufexis, & Sanchez, 2020). The animals were transported from the ENPRC Field Station to the Imaging Center with their mothers during infancy (2 weeks and 6 months) and by themselves when they were older (16 months and as adults). All subjects were scanned in a standardized supine position, using a custom-made head holder with ear bars and a mouthpiece. To facilitate laterality determination, a vitamin E capsule was taped to the right temple of each animal. Following the scanning session and full recovery of the animals, infants were immediately reunited with their mothers in shared housing. All animals were returned to their respective social groups on the following day.

T1-weighted scans were acquired using a 3D magnetization prepared rapid gradient echo (3D-MPRAGE) parallel imaging sequence (TR/TE = 2600/3.46msec -except for adults, where TR/TE 2600/3.38msec-, voxel size: 0.5mm^3^ isotropic, 8 averages, GRAPPA, R=2). T2-weighted scans were collected in the same direction as the T1-weighted scans (TR/TE = 3200/373msec, voxel size: 0.5mm^3^ isotropic, 3 averages, GRAPPA, R=2), and were combined with the T1-weighted images in the AutoSeg processing pipeline to optimize brain-atlas registration and enhance tissue segmentation (Knickmeyer et al., 2010).

Scans were collected under isoflurane anesthesia (1% to effect, inhalation) following induction with telazol (2-4 mg/kg body weight, i.m.) and intratracheal intubation to prevent motion artifacts. All animals were continuously monitored throughout the scan with an oximeter, electrocardiograph, rectal thermometer, and blood pressure monitor, while maintaining hydration dextrose/NaCl (0.45%) administered through an I.V. catheter, and stabilizing body temperature with an MRI-compatible heating pad.

#### 2.3.2 sMRI Data Processing

The structural MRI data was processed using two open-source, C++ based pipelines developed by the Neuro Image Research and Analysis Laboratories of the University of North Carolina, AutoSeg (version 3.3.2) and NeoSeg (version 1.0.7) (Styner et al., 2007; Wang et al., 2014). These semi-automatic pipelines were used to segment T1- and T2-weighted brain images into probabilistic tissue maps of gray matter (GM), white matter (WM), and cerebrospinal fluid (CSF), and generate parcellations of cortical lobes and subcortical regions of interest (ROIs) for volumetric analyses as well as total intracranial volume (ICV) as a proxy for brain size following published protocols for the rhesus brain by our group (Godfrey et al., 2023; Kovacs-Balint et al., 2021; Kovacs-Balint et al., 2023; Reding et al., 2020). NeoSeg was used combined with AutoSeg to process neonatal brain MRI images (only at 2 weeks) with intrinsic challenges such as unmyelinated WM and GM/WM contrast inversion (Prastawa, Gilmore, Lin, & Gerig, 2005).

MRI image processing steps included: 1) Averaging the T1 and the T2 images -respectively-; 2) Intensity inhomogeneity correction; 3) Rigid body registration of the images to reference atlas space; 4) Tissue segmentation and skull-stripping using atlas-based classification (ABC); 5) Registration of the atlas to the subject’s brain to generate cortical parcellations (affine + deformable ANTS registration) and computations of the respective volumes by hemisphere. For this study we used age-specific T1- and T2-weighted atlases of the infant rhesus brain generated by our group (Y Shi et al., 2017); based on best match of neuroanatomical characteristics, we registered the 2-week scans to a 2-week atlas, the 6-month scans to a 6-month atlas, and the 16-months and 7-yrs scans to a 12-months atlas. The age-specific T1- and T2-weighted atlases were employed for multi-modal registration of the subject’s brain to reference atlases for the purpose of ROI parcellation, single-atlas structural segmentation and tissue classification. At each time point, both T1- and T2-weighted data from each subject were employed jointly in full multi-modal fashion for all registration and segmentation steps of data processing, and thus both are considered of equal importance. Particularly at the youngest ages, the inclusion of the T2-weighted data is crucial for good segmentation results. Cortical ROIs were generated and defined based on published macaque MRI anatomical lobar parcellations (Knickmeyer et al., 2010) mapped onto the infant macaque brain atlases (Y Shi et al., 2017) and manually edited to ensure accurate anatomical definition following published criteria (Reding et al., 2020; Saleem & Logothetis, 2012). Developmental changes in cortical lobar volumes were analyzed as raw ROI volumetric data, as well as corrected by ICV (entered as a covariate in the statistical models) to account for individual differences in whole brain volume at each age.

#### 2.3.3 Regions of Interest (ROIs)

**Fig.2** displays the parcellations of cortical and subcortical regions studied here, which were defined based on neurohistological, cytoarchitectonic and connectivity/functional criteria (Barbas & Pandya, 1989; Carmichael & Price, 1994; Evrard, 2019) as well as published macaque MRI anatomical parcellations (Paxinos, Huang, & Toga, 2000) and previous publications by our group (Godfrey et al., 2023; Kovacs-Balint et al., 2021; Kovacs-Balint et al., 2023; Reding et al., 2020).

**Figure 2.**
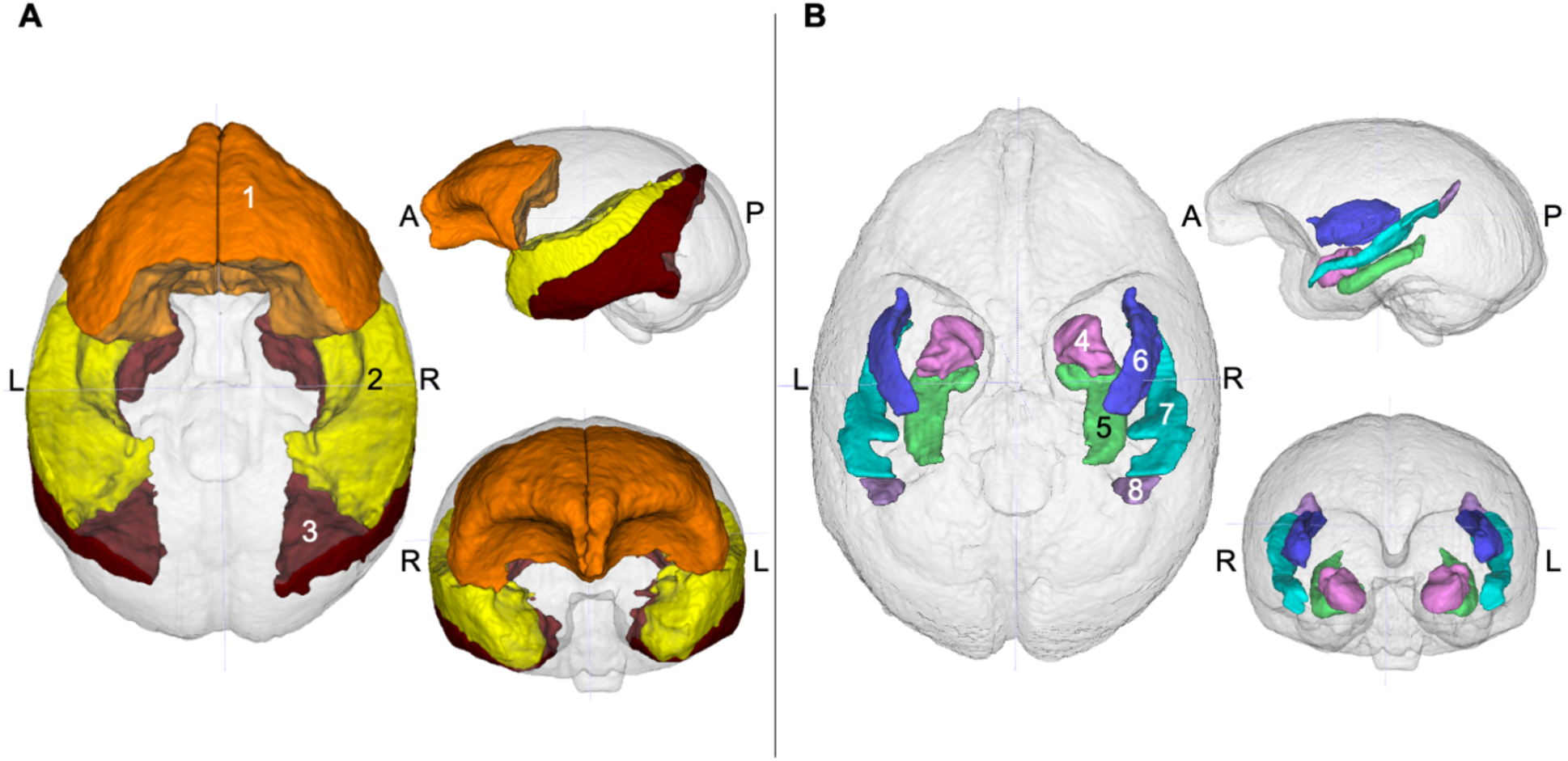
Cortical and subcortical regions of interests (ROIs). ROIs were defined based on neurohistological, cytoarchitectonic and connectivity/functional criteria. **A)** Cortical ROIs: 1) PFC cortex (brown); **B)** Subcortical ROIs and the superior temporal sulcus (STS, defined as TPOr + TPOc): 4) Amygdala (pink), 5) Hippocampus (green), 6) Insula (dark blue), 7) TPOr (turquoise), 8) TPOc (purple).

ICV was defined as total GM + total WM + total CSF in the ventricles and in the subarachnoid cavity, and volumes of each cortical lobe ROI (PFC, temporal auditory, temporal visual) were defined as the total GM + total WM (Knickmeyer et al., 2010; Kovacs-Balint et al., 2023; Reding et al., 2020; Y Shi et al., 2017; Styner et al., 2007). The total insular (INS) volume was defined as the agranular insula -AI- + dysgranular insula -DI- + granular insula -GI- + proisocortex -IPro- following rhesus monkey insula anatomical definitions (Barbas & Pandya, 1989; Carmichael & Price, 1994; Evrard, 2019); see Figs. 2B and 6. The superior temporal sulcus (STS) volume was defined and analyzed as TPOr + TPOc (Figs. 2B, 7 and 8), following anatomical criteria published for macaques (Paxinos et al., 2000; Benjamin Seltzer & Pandya, 1978). ROIs were first mapped onto the juvenile (12 months) macaque brain atlas (Shi et al, 2017) and manually edited based on the macaque specific neuroanatomical landmarks described above and following guidance from an expert rhesus neuroanatomist in our group. They were then backpropagated to the younger atlases, where the process was repeated.

### 2.4 Statistical Analysis

Analyses were conducted using IBM SPSS Statistics v.28.0.1.0 software (IBM Corporation, Armonk, NY). For each statistical model, p<0.05 was considered significant, and effect sizes were calculated as η_p_^2^. Variables were summarized as mean±standard error of the mean (SEM) to visualize trends in figures. Statistical models assumed that the data is normally distributed; thus, as a first step normality assumptions were assessed using the Shapiro-Wilk test at each age point (2weeks, 6months, 16months, Adulthood). Data that violated that assumption at 2 or more ages were log10-transformed. In addition to the results of the Shapiro-Wilk test, data histograms were inspected to assess normal distribution of the data, and outliers in the third interquartile range (IQR3) were marked. If log10-transformation did not improve the distribution of the histogram, the non-transformed data was utilized in the statistical model.

Linear Mixed Model (LMM) analysis was utilized with Age (2 weeks, 6 months, 16 months, adulthood (~7yrs)) as a repeated and fixed factor, and social Rank (DOM, SUB) and Diet (LCD-only, Choice) as fixed factors. To account for differences in overall brain size, all statistical analyses were run with and w/o ICV included as a covariate (random effect). ICV was log-transformed - and when adding ICV as a covariate to the models, brain region (at every age) were log-transformed too (even if log-transformation was not necessary for LMM).

## 3. Results

### 3.1 Intracranial Volume (ICV)

Log-transformed ICV data passed the Shapiro-Wilk test of normality, except for the 6-month and adult timepoints – therefore ICV data was log-transformed. The LMM detected a main effect of AGE (F(3, 41.432)=269.242, p=3.433×10^−27^), RANK (F(1, 87.475)=5.545, p=0.021), and DIET (F(1, 87.475)=9.512, p= 0.003) (Fig.3). Post hoc pairwise comparisons revealed that ICV volume significantly increased from 2 weeks to 6 months (p=1.317×10x^−25^), and plateaued after that (6-16mo: N.S., 16mo-7yrs: N.S.). Subordinate subjects had bigger total ICV volumes compared to DOM ones, and subjects on Choice diet also developed bigger total brain volume than subjects on low-calorie diet.

**Figure 3.**
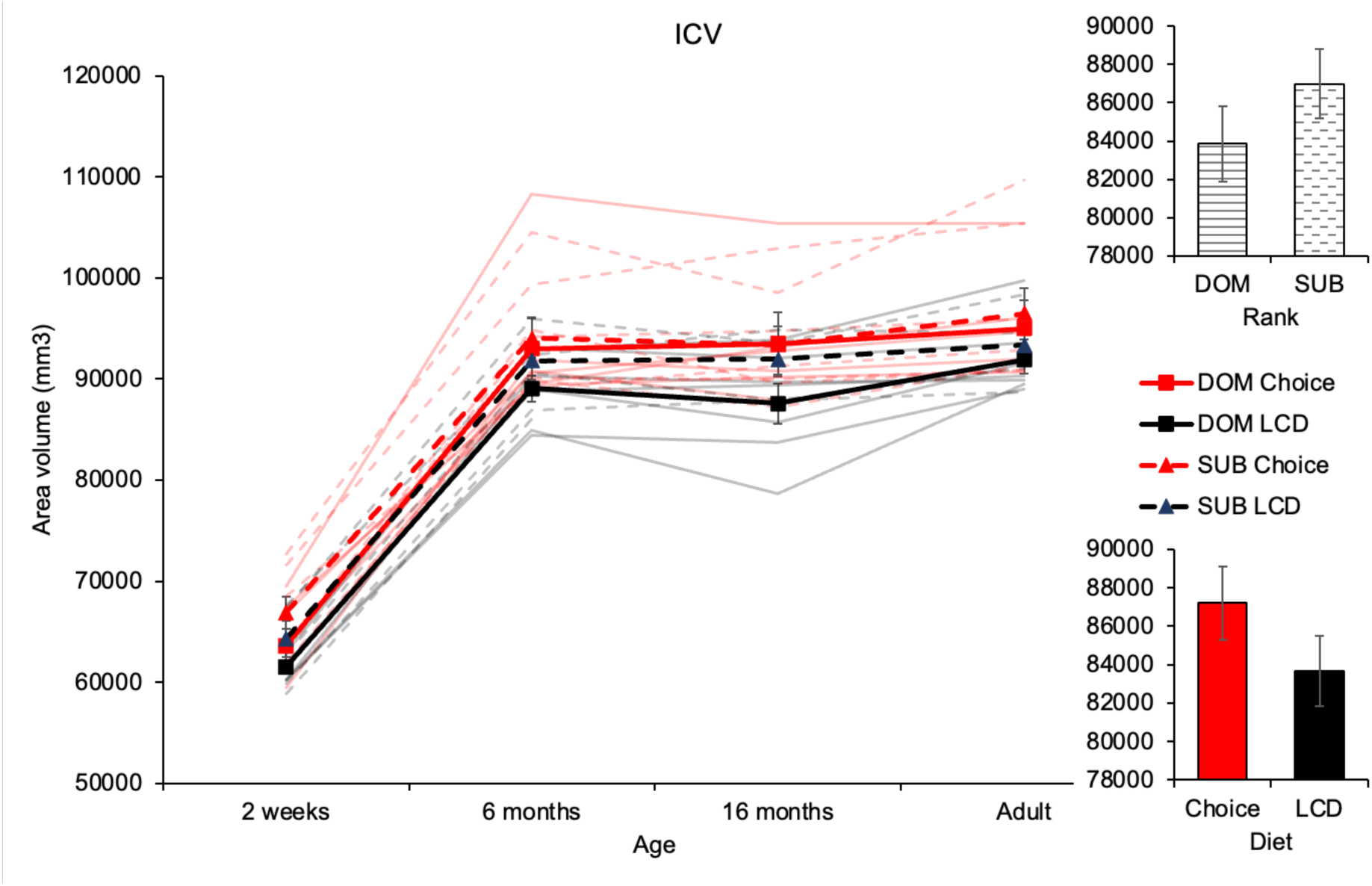
Effects of rank and diet on ICV development. Statistical analysis revealed that ICV volume increased from 2wk to 6mo of age (p=5.78×10x^−23^), and plateaued after that. Subordinate subjects showed higher ICV volumes compared to DOM subjects (p=0.021), and subjects on Choice diet showed higher total ICV than subjects on postnatal LCD (p=0.003). Thick red and black (continuous and dashed) lines represent group mean (± SEM) of ICV, while thinner and more opaque lines represent the individual ICV development by age.

### 3.2 Prefrontal Cortex (PFC)

Total PFC volume (Fig.4) data passed the Shapiro-Wilk test of normality, except at the adult timepoint. The LMM detected a main effect of AGE (F(3, 41.685)=155.353, p=1.185×10^−22^) and DIET (F(1, 78.015)=14.125, p=3.285×10^−4^). Post Hoc pairwise comparisons revealed that PFC volume increased from 2 weeks to 16 months of age (2wk-6mo: p=2.723×10^−18^, 6mo-16mo: p=0.025), then decreased by adulthood (16mo-7yr: p=0.020). Subjects on Choice diet had greater PFC volume compared to subjects on LCD (p=3.285×10^−4^).

**Figure 4.**
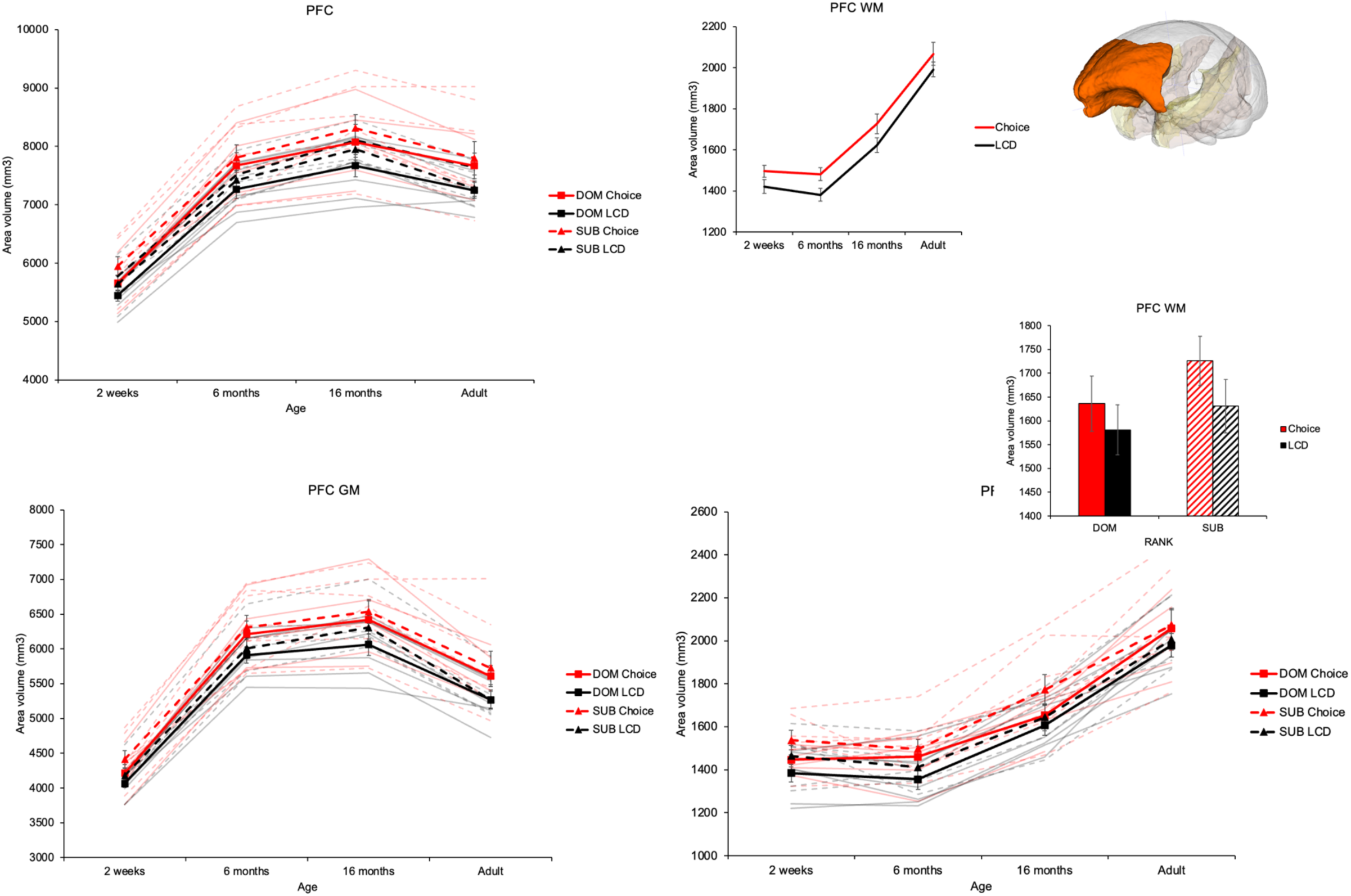
Effects of rank and diet on PFC development. Total PFC volume increased until the juvenile period (2wk-6mo: p=2.723×10^−18^, 6mo-16mo: p=0.025), and started decreasing after that (16mo-7yr: p=0.020). Prefrontal WM volume continuously increased after infancy (6-16mo: p=1.951×10^−6^, 16mo-7yr: p=9.804×10^−9^); while PFC GM volume increased during infancy (2wk-6mo: p=6.610×10^−21^), plateaued after that (6mo-16mo: p=N.S.), and started decreasing around adulthood (16mo-7yr: p=0.040×10^−7^). Thick red and black (continuous and dashed) lines represent group mean (± SEM) of ICV, while thinner and more opaque lines represent the individual ICV development by age. Total PFC volume and prefrontal GM volume changed in parallel with ICV growth. Postnatal DIET affected the PFC WM changes; SUB subjects on Choice postnatal diet had greater PFC WM volume compared to DOMs.

When ICV was added as a covariate, it had a significant effect (F(1, 57.923)=63.940, p=6.270×10^−11^). The main effect of AGE and DIET disappeared, and no other main or interaction effects were detected.

#### Prefrontal Gray Matter (GM)

Prefrontal GM volume (Fig.4) data passed the Shapiro-Wilk test of normality, except at the adult timepoint. The LMM detected a main effect of AGE (F(3, 42.716)=198.350, p=4.3123×10^−25^) and DIET (F(1, 76.788.)=12.562, p=6.740×10^−4^). Post Hoc pairwise comparisons revealed that PFC GM volume increased from 2 weeks to 6 months of age (p=6.610×10^−21^), plateaued after that (6mo-16mo: p=N.S.), and started decreasing around adulthood (16mo-7yr: p=0.040×10^−7^). Subjects on Choice diet had greater PFC volume compared to subjects on LCD (p=3.285×10^−4^).

When ICV was added as a covariate, significant effect of total ICV (F(1, 71.046)=31.836, p=3.198×10^−7^) was observed. The main effect of AGE and DIET disappeared, and no other main or interaction effects were detected.

#### Prefrontal White Matter (WM)

Prefrontal WM volume (Fig.4) data passed the Shapiro-Wilk test of normality. The LMM detected a main effect of AGE (F(3, 36.490)=81.955, p=2.821×10^−16^), RANK (F(1, 74.191)=4.226, p=0.043) and DIET (F(1, 74.191)=8.307, p=0.005). Post Hoc pairwise comparisons revealed that PFC WM volume did not change early on (2wk-6mo: N.S.), but started increasing after that (6-16mo: p=1.951×10^−6^, 16mo-7yr: p=9.804×10^−9^). Subordinate subjects showed higher WM volume compared to DOMs (p=0.043); and subjects on Choice infant diet showed greater WM volume compared to LCD subjects (p=0.005).

When ICV was added as a covariate, significant effect of total ICV (F(1, 57.764)=22.541, p=1.398×10^−5^) was observed. The main effect of AGE, DIET and RANK disappeared, but significant AGE x DIET (F(3, 23.576)=6.550, p=0.002), DIET x RANK (F(1, 58.060)=12.408, p=8.4×10^−4^), AGE x DIET x ICV (F(3, 23.460)=6.600, p=0.002) and DIET x RANK x ICV (F(1, 57.764)=12.400, p=8.456×10^−4^) interaction effects were detected.

### 3.3 Temporal Visual Area (TVA)

The temporal visual area volume (Fig.5) data passed the Shapiro-Wilk test of normality, except at the 2-week timepoint. The LMM detected a main effect of AGE (F(3, 41.639)=175.693, p=1.162×10^−23^) and RANK (F(1, 82.639)=8.044, p=0.006). Post Hoc pairwise comparisons revealed that the volume of the temporal visual area increased early on (2wk-6mo: p=1.907×10^−18^), and plateaued after that (6-16mo: p=N.S., 16mo-7yr: p=N.S.). Subordinate subjects had greater temporal visual lobe volume compared to DOMs (p=0.006).

**Figure 5.**
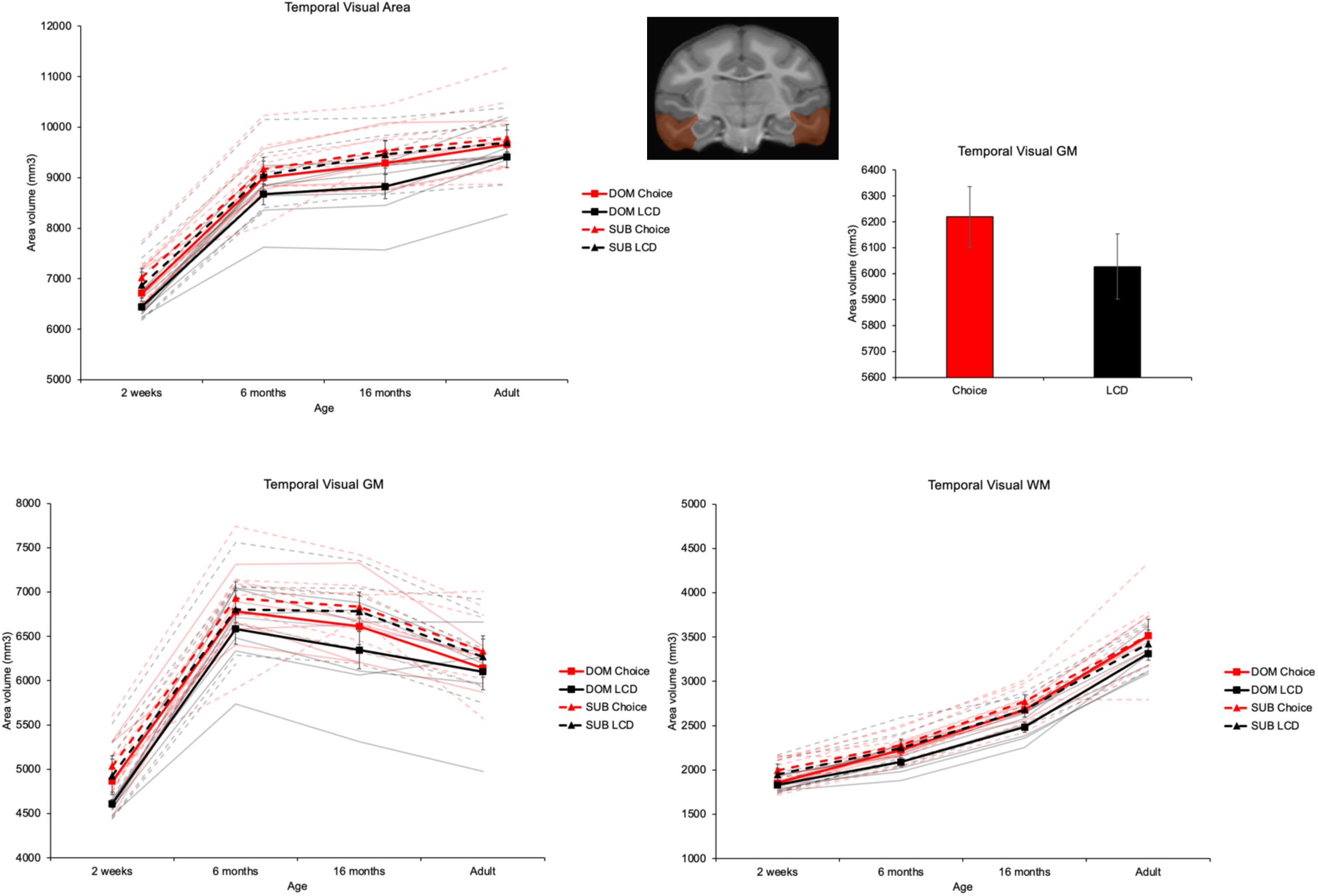
Effects of rank on temporal visual area (TVA) development. Total TVA volume increased early on (2wk-6mo: p=1.907×10^−18^), and plateaued after that (6-16mo: p=N.S., 16mo-7yr: p=N.S.). TVA WM volume increased continuously until adulthood (2wk-6mo: p=6.605×10^−8^; 6-16mo: p=1.977×10^−10^; 16mo-7yr: p=1.918×10^−11^); while TVA GM volume increased during infancy (2wk-6mo: p=1.249×10^−19^), and started decreasing around the juvenile period (16mo-7yr: p=0.040×10^−7^). Thick red and black (continuous and dashed) lines represent group mean (± SEM) of ICV, while thinner and more opaque lines represent the individual ICV development by age. After controlling for the total ICV, subjects on Choice postnatal diet showed greater TVA GM volume compared to subjects on LCD.

When ICV was added as a covariate, it had a significant effect (F(1, 41.994)=160.196, p=6.366×10^−16^). The main effect of AGE, RANK and DIET disappeared, and no other main or interaction effects were detected.

#### Temporal Visual Area Gray Matter (TVA GM)

The TVA GM volume (Fig.5) data passed the Shapiro-Wilk test of normality, except at the 2-week timepoint. The LMM detected a main effect of AGE (F(3, 43.072)=140.310, p=2.979×10^−22^) and RANK (F(1, 82.217)=7.183, p=0.009). Post Hoc pairwise comparisons revealed that TVA GM volume sharply increased from 2 weeks to 6 months (p=1.249×10^−19^), and started decreasing during the juvenile period (16mo-7yrs: p=0.014). Subordinate subjects showed greater TVA GM volume compared to DOMs (p=0.009).

When ICV was added as a covariate, significant effect of total ICV (F(1, 42.236)=144.360, p=3.263×10^−15^) was observed. The main effect of AGE and RANK disappeared, while main effect of DIET (F(1, 42.384)=8.163, p=0.007), and interaction effect of DIET x ICV (F(1, 42.236)=8.110, p=0.007) emerged. Post Hoc pairwise comparisons revealed that subjects on Choice diet developed greater TVA GM volume compared to subjects on LCD postnatally.

#### Temporal Visual Area White Matter (TVA WM)

The TVA WM volume (Fig. 5) data passed the Shapiro-Wilk test of normality, except at the 2-week timepoint. The LMM detected a main effect of AGE (F(3, 34.704)=193.419, p=1.0137×10^−21^). Post Hoc pairwise comparisons revealed that TVA WM volume increased continuously until adulthood (2wk-6mo: p=6.605×10^−8^; 6-16mo: p=1.977×10^−10^; 16mo-7yr: p=1.918×10^−11^).

When ICV was added as a covariate, significant effect of total ICV (F(1, 55.898)=28.144, p=1.996×10^−6^) was observed. The main effect of AGE disappeared, and no other main or interaction effects were detected.

### 3.4 Temporal Auditory Area (TAA)

The temporal auditory area volume (Fig. 6) data passed the Shapiro-Wilk test of normality. The LMM detected a main effect of AGE (F(3, 41.832)=225.910, p=7.325×10^−26^), RANK (F(1, 79.391)=8.736, p=0.004), and DIET (F(1, 79.391)=6.968, p=0.01). Post Hoc pairwise comparisons revealed that TAA volume increased until the juvenile period (2wk-6mo: p=3.281×10^−19^; 6-16mo: p=4.860×10^−4^), and plateaued after that (16mo-7yr: p=N.S.). Subordinate subjects showed higher TAA volume than DOMs (p=0.004), and subjects on Choice postnatal diet developed greater TAA volume compared to subjects on LCD (p=0.01).

**Figure 6.**
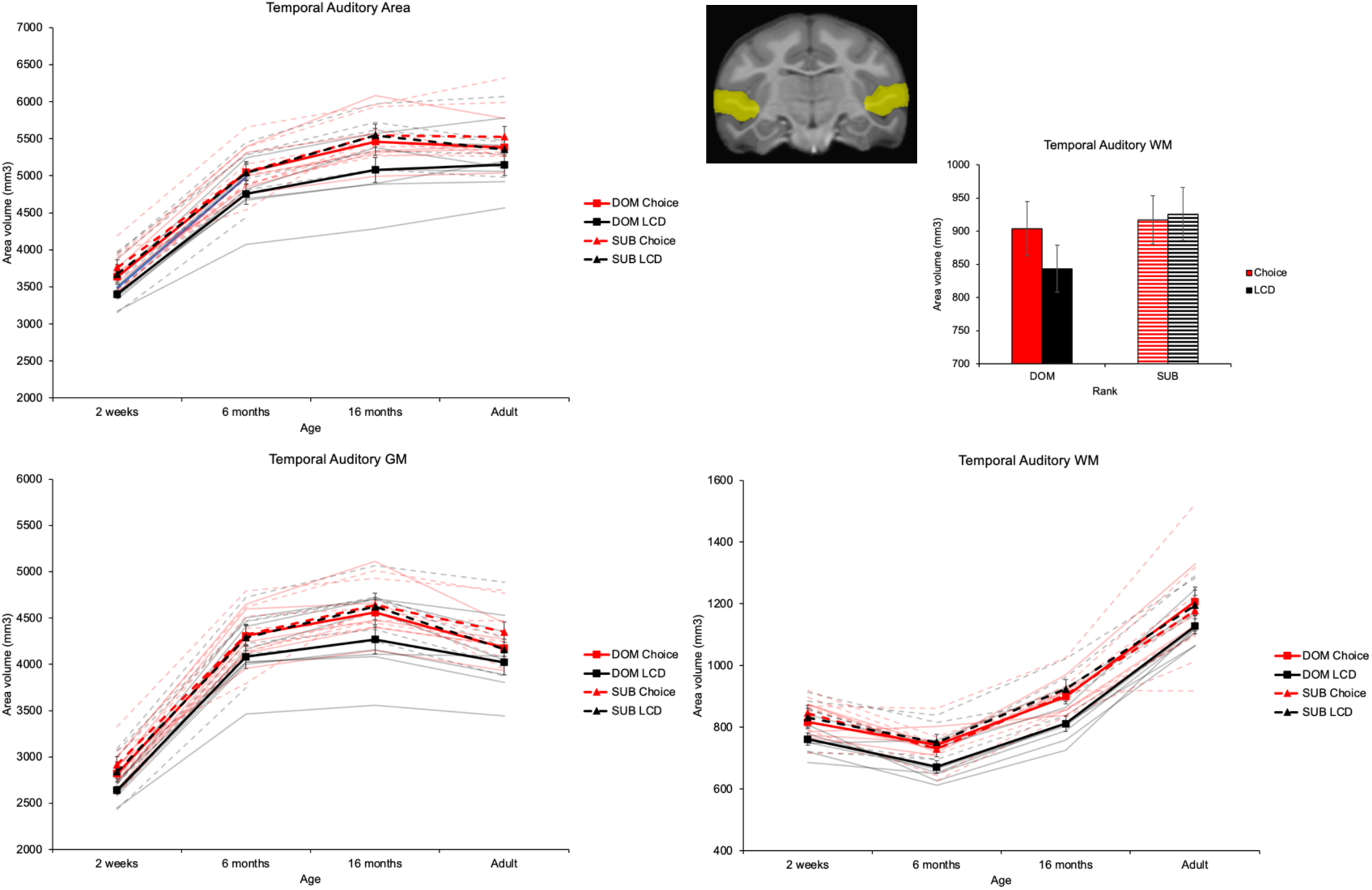
Effects of rank and diet on temporal auditory area (TAA) development. Total TAA volume increased until the juvenile period (2wk-6mo: p=3.281×10^−19^; 6-16mo: p=4.860×10^−4^), and plateaued after that (16mo-7yr: p=N.S.). TAA GM volume increased until the juvenile period (2wk-6mo: p=6.625×10^−21^; 6-16mo: p=0.029), and started decreasing before adulthood (16mo-7yr: p=0.004). TAA WM volume significantly decreased after birth / during infancy (2wk-6mo: p=1.640×10^−5^), and steadily increased after that (6-16mo: p=3.657×10^−11^; 16mo-7yr: p=3.484×10^−11^). Thick red and black (continuous and dashed) lines represent group mean (± SEM) of ICV, while thinner and more opaque lines represent the individual ICV development by age.

When ICV was added as a covariate, significant effect of total ICV (F(1, 67.723)=121.534, p=9.223×10^−17^) was observed. The main effect of AGE, DIET and RANK disappeared, and no other interaction effects were detected.

#### Temporal Auditory Area Gray Matter (TAA GM)

The TAA GM volume (Fig. 6) data passed the Shapiro-Wilk test of normality. The LMM detected a main effect of AGE (F(3, 43.851)=265.218, p=4.071×10^−28^), RANK (F(1, 77.995)=6.883, p=0.01), and DIET (F(1, 77.995)=5.888, p=0.018). Post Hoc pairwise comparisons revealed that TAA GM volume continuously increased from birth until the juvenile period (2wk-6mo: p=6.625×10^−21^; 6-16mo: p=0.029), and started decreasing before adulthood (16mo-7yr: p=0.004). Subordinate subjects showed higher TAA GM volume compared to DOMs (p=0.01), and subjects on Choice postnatal diet developed greater TAA volume compared to subjects on LCD (p=0.018).

When ICV was added as a covariate, significant effect of total ICV (F(1, 56.680)=80.393, p=1.84×10^−12^) was observed. The main effect of AGE, DIET and RANK disappeared, and no other interaction effects were detected.

#### Temporal Auditory Area White Matter (TAA WM)

The TAA WM volume (Fig. 6) data passed the Shapiro-Wilk test of normality. The LMM detected a main effect of AGE (F(3, 35.511)=100.454, p=2.062×10^−17^) and RANK (F(1, 55.443)=6.161, p=0.016), and a significant RANK x DIET interaction (F(1, 55.443)=6.620, p=0.013). Post Hoc pairwise comparisons revealed that TAA WM volume significantly decreased after birth / during infancy (2wk-6mo: p=1.640×10^−5^), and steadily increased after that (6-16mo: p=3.657×10^−11^; 16mo-7yr: p=3.484×10^−11^). Subordinate subjects showed greater TAA WM volume compared to DOMs (p=0.016). Subordinate subjects on postnatal LCD showed greater TAA WM volume compared to DOM subjects on LCD.

When ICV was added as a covariate, significant effect of total ICV (F(1, 26.529)=14.521, p=7.452×10^−4^) was observed. No other main or interaction effects were detected.

### 3.5 Superior Temporal Sulcus (STS)

#### Temporal Parieto-Occipital Area, Rostral Part (TPOr)

The TPOr volume (Fig.7) data passed the Shapiro-Wilk test of normality. The LMM detected a main effect of AGE (F(3, 38.253)=354.169, p=6.143×10^−28^), RANK (F(1, 65.032)=11.734, p=0.001), and DIET (F(1, 65.032)=6.977, p=0.010). Post Hoc pairwise comparisons revealed that the TPOr volume increased steadily from birth until the juvenile period (2wk-6mo: p=2.443×10^−14^; 6-16mo: p=4.731×10^−18^), and started decreasing after that (16mo-7yr: p=5.559×10^−6^). Subordinate subjects showed greater TPOr volume compared to DOMs (p=0.001), and subjects on Choice postnatal diet developed greater TPOr volume compared to subjects on LCD (p=0.010).

**Figure 7.**
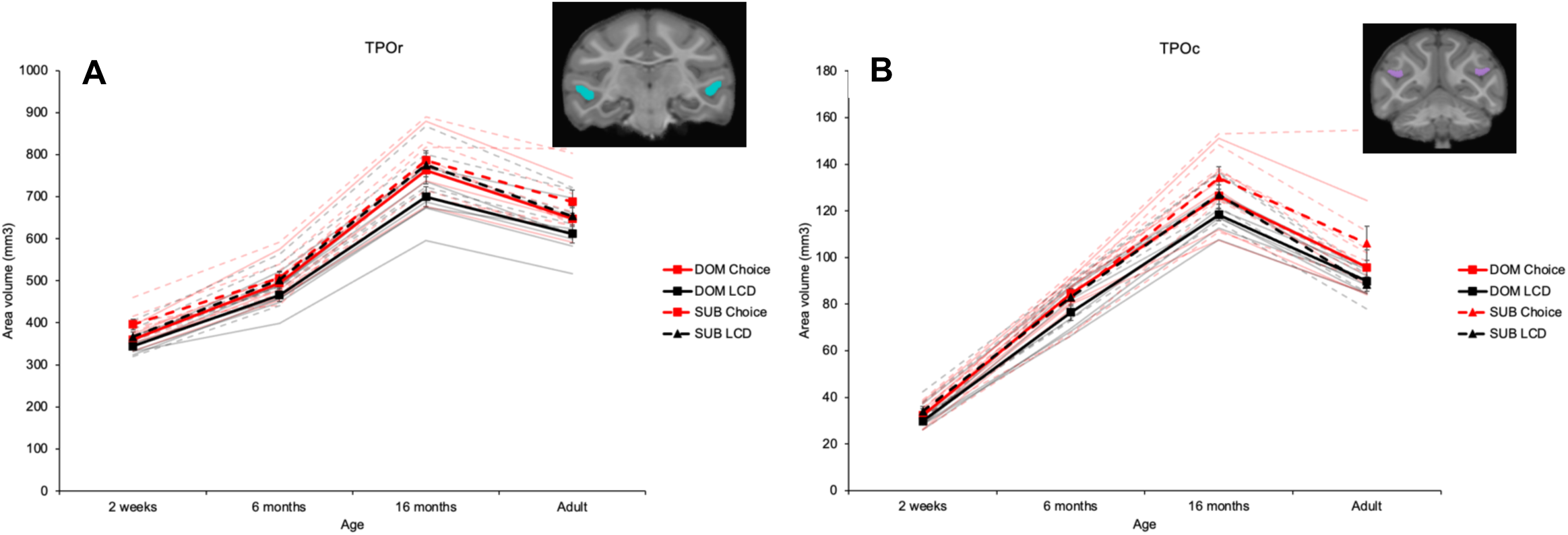
Effects of rank and diet on Superior Temporal Sulcus (STS) development. Both the rostral and the caudal part of the TPO grew steadily from birth until the juvenile period (TPOr: 2wk-6mo: p=2.443×10^−14^, 6-16mo: p=4.731×10^−18^; TPOc: 2wk-6mo: p=1.587×10^−28^, 6-16mo: p=2.452×10^−18^), and started decreasing after that (TPOr: 16mo-7yr: p=5.559×10^−6^; TPOc: 16mo-7yr: p=1.724×10^−10^). Thick red and black (continuous and dashed) lines represent group mean (± SEM) of ICV, while thinner and more opaque lines represent the individual ICV development by age.

When ICV was added as a covariate, significant effect of total ICV (F(1, 55.870)=63.907, p=8.028×10^−11^) was observed. No other main or interaction effects were detected.

#### Temporal Parieto-Occipital Area, Caudal Part (TPOc)

TPOc volume (Fig.7B) data passed the Shapiro-Wilk test of normality, except at the 6 month and adult timepoints – therefore log-transformed data was used at each timepoint. The LMM detected a main effect of AGE (F(3, 39.466)=617.342, p= 3.476×10^−33^), RANK (F(1, 83.912)=5.787, p= 0.018) and DIET (F(1, 83.912)=7.476, p=0.008). Post Hoc pairwise comparisons revealed that the TPOc volume increased steadily from birth until the juvenile period (2wk-6mo: p=1.587×10^−28^; 6-16mo: p=2.452×10^−18^), and started decreasing after that (16mo-7yr: p=1.724×10^−10^). Subordinate subjects showed greater TPOc volume compared to DOMs (p=0.018), and subjects on Choice postnatal diet developed greater TPOc volume compared to subjects on LCD (p=0.008).

When ICV was added as a covariate, no significant effect of total ICV was observed. No other main or interaction effects were detected.

### 3.6 Insula

Insula volume (Fig. 8A) data passed the Shapiro-Wilk test of normality. The LMM detected a main effect of AGE (F(3, 48.137)=158.959, p=5.678×10^−25^), RANK (F(1, 75.651)=6.695, p=0.012), and DIET (F(1, 75.651)=18.262, p=5.542×10^−5^). Post Hoc pairwise comparisons revealed that the insula volume increased during infancy (2wk-6mo: p=2.472×10^−16^), plateaued during the juvenile period (6-16mo: p=N.S.), and started decreasing after that (16mo-7yr: p=0.018). Subordinate subjects showed greater insula volume than DOMs (p=0.012), and subjects on Choice postnatal diet showed greater insula volume compared to subjects on LCD (p=5.542×10^−5^).

**Figure 8.**
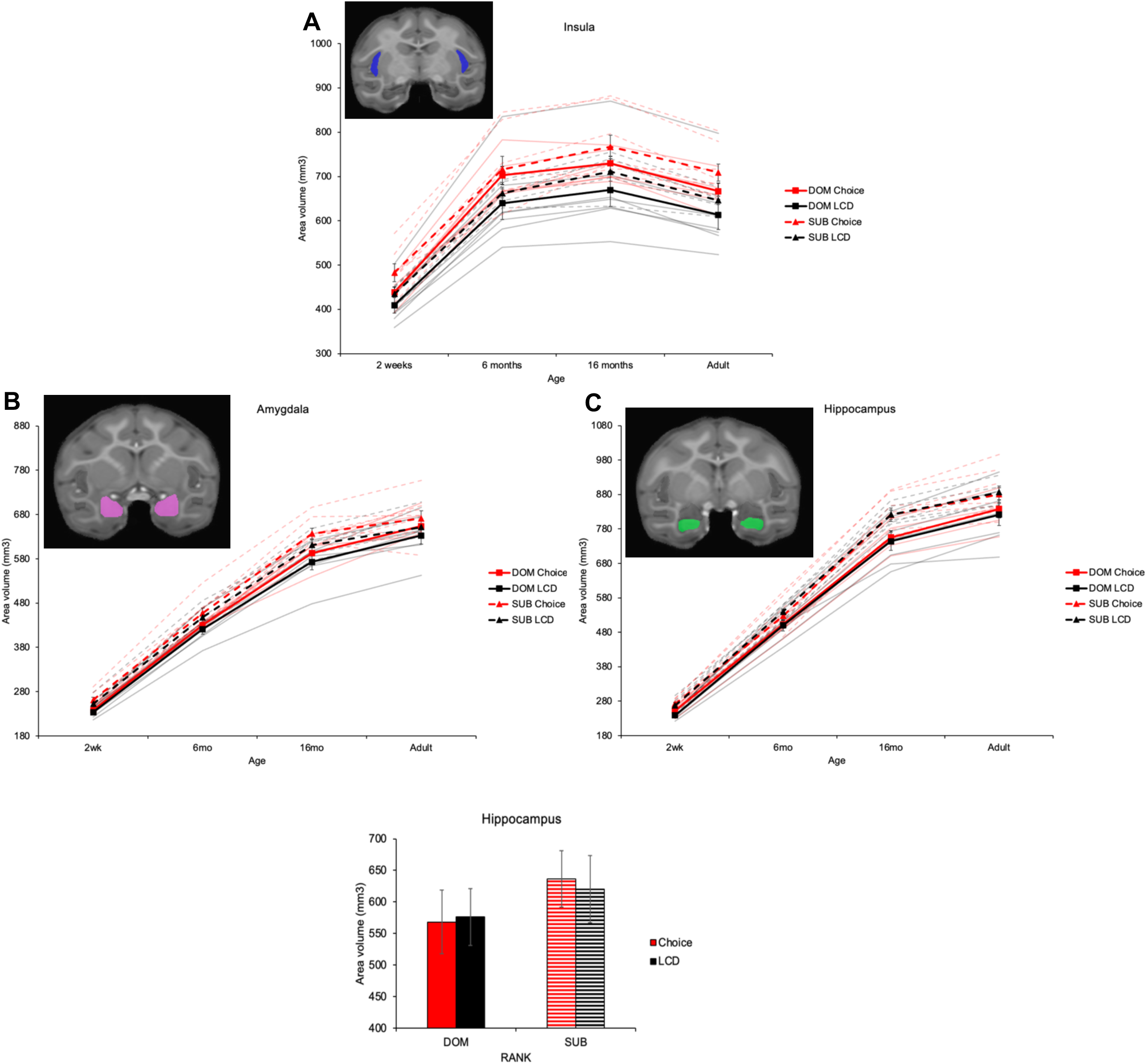
Effects of rank and diet on Subcortical development. **A)** Insula volume increased during infancy (2wk-6mo: p=2.472×10^−16^), plateaued during the juvenile period (6-16mo: p=N.S.), and started decreasing after that (16mo-7yr: p=0.018). **B)** Amygdala volume continuously increases from birth to adulthood (2wk-6mo: p=1.665×10^−26^, 6-16mo: p=4.183×10^−19^, 16mo-7yr: p=9.413×10^−4^). **C)** Hippocampus volume rapidly increased from birth through the juvenile period, and continued to slowly increase through adulthood (2wk-6mo: p=1.096×10^−24^, 6-16mo: p=4.857×10^−21^, 16mo-7yr: p=8.006×10^−4^). When controlling for the individual brain size, DIET x RANK interaction suggested greater hippocampus volume in SUB subjects on LCD compared to Choice diet. SUB subjects on LCD developed greater hippocampus than DOMs.

When ICV was added as a covariate, significant effect of total ICV (F(1, 66.542)=25.430, p=3.768×10^−6^) was observed. No other main or interaction effects were detected.

### 3.7 Amygdala

Amygdala volume (Fig. 8B) data violated the Shapiro-Wilk test of normality only at 6months of age – therefore data was not transformed for any of the ages. The LMM detected a main effect of AGE (F(3, 36.756)=1313.492, p=2.021×10^−37^), RANK (F(1, 61.483)=16.465, p=1.422×10^−4^) and DIET (F(1, 61.483)=5.020, p=0.029). Post Hoc pairwise comparisons revealed that the AMY volume continuously increases from birth to adulthood (2wk-6mo: p=1.665×10^−26^, 6-16mo: p=4.183×10^−19^, 16mo-7yr: p=9.413×10^−4^). Subordinate subjects showed greater AMY volume than DOMs (p=1.422×10^−4^), and subjects on Choice postnatal diet showed greater AMY volume compared to subjects on LCD (p=0.029).

When ICV was added as a covariate, it had a significant effect in the model (F(1, 61.652)=30.987, p=6.011×10^−7^). The main effect of AGE, RANK and DIET disappeared, and no other interaction effects were detected.

### 3.8 Hippocampus

Hippocampus volume (Fig.8C) data passed the Shapiro-Wilk test of normality. The LMM detected a main effect of AGE (F(3, 35.551)=1544.084, p=1.076×10^−37^) and RANK (F(1, 58.104)=23.384, p=1.012×10^−5^). Post hoc pairwise comparisons revealed that the hippocampus volume rapidly increased from birth through the juvenile period, and continued to slowly increase through adulthood (2wk-6mo: p=1.096×10^−24^, 6-16mo: p=4.857×10^−21^, 16mo-7yr: p=8.006×10^−4^). Subordinate subjects showed greater hippocampus volume than DOMs (p=1.012×10^−5^).

When ICV was added as a covariate, it had a significant main effect (F(1, 59.954)=8.041, p=0.006) and DIET x RANK x ICV interaction effect in the model (F(1, 59.954)=9.363, p=003). Although the main AGE and RANK effects disappeared, a significant DIET x RANK effect was detected (F(1, 60.037)=9.487, p=0.003). The results shown in Figure 8C suggest that this interaction effect might be driven by bigger hippocampal volume in SUB than DOM animals exposed to the postnatal LCD diet (p=0.003).

## 4. Discussion

The goal of this study was to examine the developmental and long-term effects of postnatal exposure to social subordination stress and obesogenic diets on the structure of brain regions involved in stress and emotional regulation, social processing, and homeostasis. Brain structural MRI scans were collected longitudinally from a cohort of socially-housed female rhesus macaques with extreme social status differences (DOM, SUB), and diet exposures (LCD, Choice) from infancy through the juvenile period and again as young adults. After menarche, all subjects were maintained on an LCD-only diet through adulthood. Our findings show insult-specific effects of obesogenic diet and psychosocial stress on cortical and corticolimbic brain regions. Animals with access to the obesogenic diet had larger overall brain size (measured as ICV) and larger PFC, PFC GM, INS, TPOr, and TPOc than those in the LCD-diet only; most of these regional diet effects, expect for the INS, were driven by general effects of the diet on brain size. The diet effects were lost when adding the adult data to the longitudinal analysis, suggesting transient effects of obesogenic diets while the animals were consuming it, but not long-term, persistent effects. These findings highlight the potential of brain rescue mechanisms that could offset lasting developmental effects of early-life obesogenic diet consumption. With respect to social rank, region-specific effects were detected, with larger volumes of hippocampus, TPOr and temporal auditory cortices in SUB than DOM animals. These findings confirmed the initial hypothesis of enlarged brain regions involved in socioemotional processing and monitoring of faces (Andrews, 2005) in SUB than DOM animals, even after controlling for individual differences in brain size, suggesting long-term, persistent and cumulative effects of adverse social experiences, in contrast to the transient diet effects.

Subjects with postnatal exposure to obesogenic diets showed larger overall brain size (measured as ICV) as well as larger volumes in the PFC, PFC GM, INS, STS, TPOr, and TPOc than those in the LCD-diet only. After correcting for ICV, however, the diet effects on these ROIs were lost, with the exception of the INS, suggesting that most effects were not region-specific, but driven by the effects of diet on brain size. The diet effects were lost when adding the adult data to the longitudinal analysis, suggesting a transient effect of obesogenic diets while the animals were consuming it, but not long-term, persistent effects. These findings highlight the potential of brain rescue mechanisms that could offset lasting developmental effects of early-life obesogenic diet consumption.

Social rank had region-specific effects, with larger volumes of hippocampus, temporal visual, temporal auditory and TPOr in SUB than DOM animals. These findings confirmed the initial hypothesis of larger regions involved in social processing and monitoring of faces (Andrews, 2005) in SUB than DOM animals. The effects of social rank were still detected and prominent in STS, TPOr, and temporal auditory cortices after controlling for individual differences in brain size (ICV), and suggest persistent effects of the adverse social experience, particularly evident at later ages (16months, adulthood) suggesting a potential cumulative effect of social subordination stress long-term on the structure of brain ROIs important for social cognition and sensory processing.

A major finding in this study was the larger overall brain size (ICV) across development (longitudinally, from 2weeks to 16 months of age) in animals exposed postnatally to the obesogenic diet. This was inconsistent with previous reports of widespread GM volume reductions in overweight and obese individuals (García-García et al., 2019; Herrmann, Tesar, Beier, Berg, & Warrings, 2019; Li et al., 2022); although it should be noted that the literature on the effects of obesogenic diets on overall brain volume development is mixed. A previous study on childhood obesity in humans reported an inverse association between childhood obesity and brain GM and overall brain volumes (Jiang et al., 2022). Other studies reported larger overall brain WM and total brain volumes in obese children, instead (Shao, Tan, & He, 2022). A potential mechanism that could explain the latter findings and our results include higher energy intake from the obesogenic diets, which can lead to increased dendritic arborization and/or myelination and subsequent larger brain volumes (Yoon et al., 2016). Interestingly, the effects of the obesogenic diet were transient, with no long-term, persistent effects after the animals were put back in the healthy, LCD-only diet. Emerging research suggests that a HCD during critical periods of brain development may contribute to increased neuroinflammation and oxidative stress in the brain, which can potentially result in neuronal damage and alterations in brain structure and function (Cavaliere et al., 2019). Hence, it is plausible that the cessation of HCD consumption may have provided a restorative period for the brain to stop ongoing effects on dendritic arborization and/or myelination, and neuroinflammation/oxidative stress, potentially mitigating the long-term negative impact on brain development. Although we can only speculate about the underlying mechanisms, some could involve the impact of HCD on the gut microbiome which, in turn, could affect brain health through, for example, chronic low-grade inflammation and impairments in cognitive function (Alsegiani & Shah, 2022).

Our finding of enlarged INS in the Choice group is at odds with previous reports, as higher body mass index (BMI) and early life gains in adiposity have been linked to smaller, rather than bigger, GM volumes in the INS (Rasmussen et al., 2017). Interestingly, the INS growth rate is different in macaques than in humans, being one of the fastest-growing regions of the macaque brain during infancy (Gilmore et al., 2012; Scott et al., 2016), Given this, it may make the INS more vulnerable to diet-specific insults in macaque than human infants. Furthermore, a recent study uncovered that obesity and sweet-food addiction in females has been linked to increased INS activity (Hsu et al., 2017), thereby supporting our findings that early exposure to obesogenic diets can induce overactivity, and potentially alter INS structure, given its critical role in both feeding and stress responses (Spierling et al., 2020). Here again, as described for ICV, the diet effects were normalized by adulthood, which supported the hypothesis that cessation of an obesogenic diet may restore homeostatic balance in the body and reverse the effects on the brain (Ahmed et al., 2022).

Furthermore, long-term effects of the obesogenic diet were uncovered in the total PFC and PFC GM of these subjects. Both the developmental and long-term effects, however, were lost upon ICV-correction, indicating that these effects were not region-specific, but, rather, driven by global differences in brain size. This finding is not consistent with a large body of research demonstrating associations between increased BMI and reductions in PFC GM volumes (Opel et al., 2021); or PFC atrophy reported in obese adolescents (Bruehl, Sweat, Tirsi, Shah, & Convit, 2011; Yokum, Ng, & Stice, 2012), adults (Pannacciulli et al., 2006; Walther, Birdsill, Glisky, & Ryan, 2010), and rats fed a high fat diet (Bocarsly et al., 2015). However, our SOD subjects had not shown an obese phenotype or even increases in body weight by the time of the juvenile, prepubertal, assessments. Given the drastic remodeling that takes place in the PFC during adolescence (Casey, Jones, & Hare, 2008) and effects of obesity on pubertal development (Burt Solorzano & McCartney, 2010), it would have been interesting to maintain the animals in the obesogenic diet until late adolescence to examine potential interactions between diet and adolescence on PFC remodeling.

The lack of social rank effects on PFC were surprising. The human PFC and its subregions are particularly vulnerable to chronic psychosocial stress and early life adversity, which can lead to both physiological and volumetric alterations (Bremner, 2002, 2006). Moreover, studies have reported WM alterations in PFC regions in both children and NHPs exposed to other forms of chronic stress during development (Carrion et al., 2001; De Bellis et al., 1999; Hanson et al., 2010). The latter finding is also inconsistent with human and animal literature largely showing an association between chronic stress and smaller mPFC volumes. It should be noted, though, that PFC subregions are differentially impacted by chronic stress (Datta & Arnsten, 2019), such as mPFC shrinkage but orbitofrontal cortex (OFC) expansion in response to stress (McEwen & Morrison, 2013a). Overall, the findings regarding PFC volume may depend on the specific subregion. Hence, it would be interesting for future studies to explore which subregions are most susceptible to chronic stress-induced structural alterations to see which subregion is driving the observed effects on total PFC.

Although no diet effects were found in either TVA or TAA, significant effects of social rank were detected, with consistently bigger volumes in SUB than DOM in total TVA, TVA GM, TVA WM. Most of these effects persisted after ICV-correction, indicating that the rank effects were region-specific. The temporal visual lobe contains areas that are part of the visual ventral object pathway and play a vital role in facial and object recognition and processing of sociovisual information (Andrews, 2005; Kovacs-Balint et al., 2021). These high-level visual regions promote object discrimination and enable intentional filtering rather than local, low level, features of visual stimuli, such as contrast or orientation (Kuwahata, Adachi, Fujita, Tomonaga, & Matsuzawa, 2004; Lutz, Lockard, Gunderson, & Grant, 1998; Muschinski et al., 2016). Furthermore, given the ability of macaques to determine dominance relationships through visual observation of their peers’ behaviors (Paxton et al., 2010), the responsiveness of neurons in the temporal visual lobe may be heightened in SUB as an adaptation to pay attention and be ready to react to visual cues related to social hierarchy, thus leading to rapid dendritic spine increase and synaptic proliferation (Rakic, Bourgeois, Eckenhoff, Zecevic, & Goldman-Rakic, 1986). Visual inspection of the figures suggests that the effects of social rank on the temporal visual lobe WM are prominent as early as 2 weeks of age. The rapid development of WM in the temporal visual lobe during younger ages (Kovacs-Balint et al., 2021; Y. Shi et al., 2013) may account for the early effects of social rank on WM. This rapid increase in WM may also increase the region’s sensitivity to environmental insults during this developmental period. Additionally, studies have found that the temporal visual lobe experiences rapid growth from 2 to 16 weeks of age, followed by a plateau, further supporting these findings (Kovacs-Balint et al., 2021).

Compared to other cortical lobes, the temporal lobe undergoes significant volumetric development after 1 year of age due to its role in integrating sensory input and generating behavior (Scott et al., 2016). Furthermore, WM in the temporal cortex continues to increase during the juvenile and adolescent periods (1-5 years) (Knickmeyer et al., 2010), thereby increasing its long-term susceptibility to psychosocial stress compared to other cortical regions that serve low order functions and stabilize earlier (Scott et al., 2016). This could explain the region-specific rank effects observed. The temporal auditory lobe contains circuits responsible for auditory processing and motion perception, -such as body movement-, as well as face- and eye-movement detection (Kovacs-Balint et al., 2021). Single cell recordings have shown that certain neurons in the temporal auditory lobe are responsive to complex auditory stimuli such as monkey vocalizations and lip-smacking, which are part of submissive/appeasing behaviors displayed by SUB rhesus (Bruce, Desimone et al, 1981; Baylis, Rolls et al, 1987). Hence, from an ontogenetic and evolutionary perspective, the larger temporal visual and auditory lobe volumes in SUB may be adaptive to support increased accuracy to process and respond to social visual and auditory stimuli to navigate threats and social hierarchies throughout adulthood.

Developmental effects of diet were detected on the STS (TPOr and TPOc), with larger volumes in animals in the Choice than the LCD-only diet. These effects were lost upon correction for ICV, indicating that they were driven by global effects of the obesogenic diet on brain size. Strong effects of social rank were also detected, particularly in the TPOr up to adulthood after correcting for brain size, with SUB exhibiting larger TPOr -but not TPOc-volumes than DOM. These findings suggest larger regions involved in social processing and monitoring of faces (Andrews, 2005) across the life-span of SUB, suggesting persistent and cumulative effects of social experiences.

These findings are at odds with reports of a positive correlation between GM in the mid-STS (mSTS) of macaques and high social rank in macaques (Sallet et al., 2011). It is plausible, however, that the larger STS in SUB was driven by the high complex social environments these animals live in and their need to constantly utilize social monitoring to gather information about the location of potential aggressors and to gain insight into the emotional states and intentions of their peers (Evers, de Vries, Spruijt, & Sterck, 2012; Ramezanpour, Görner, & Thier, 2021). Although highly speculative, our findings could be compared with studies in maltreated children showing greater GM volumes in the superior temporal gyrus -STG- (De Bellis et al., 2002), potentially demonstrating the long-term adaptation of the STS to accurately read social threats.

Interestingly, a long-term effect of rank emerged in the TPOr after ICV-correction, with larger volumes in SUB. Studies have revealed that the TPO is a region within the STS that receives input from auditory, visual, and somatosensory areas (Barnes & Pandya, 1992; B. Seltzer et al., 1996) and has reciprocal connections with AMY (Amaral & Insausti, 1992). In a fMRI study of AMY connectivity, the presentation of dynamic fearful facial expressions enhanced responses in dorsal temporal areas, including STS. STS is therefore sensitive to fearful stimuli presumably via input from the AMY, which may control sensitivity of cortical areas, such as STS, to socio-emotionally relevant stimuli (Furl, Henson, Friston, & Calder, 2013). Given the tremendous amount of social threats and environmental pressures experienced by SUB, larger TPOr volumes would be advantageous to mount quick adaptive behavioral responses in complex social groups. It highlights the TPOr as a specific sub-region of the STS important for the social and emotional stimuli processing and relaying that information to areas such as the hippocampus and AMY, which in turn modulate the activity of the STS to focus on decoding stimuli relevant to fear and anxiety behaviors.

This study has several limitations. The first one is the sample size, which, although big for a study with socially-housed macaques, limits the statistical power for the analyses of important interactions between factors. It would have also been important to examine the effects of heritable/prenatal factors from the biological mother (DOM, SUB) on cross-fostered infants -or control for them as a covariate-, as well as the effects of cross-fostering, but we were underpowered to do so. Furthermore, detecting long-term significant effects of diet on adult brain structure might be limited due to the use of categorical variables (Choice, LCD) in the LMM models. An alternative statistical approach in follow up studies should include multiple regression models to examine whether total HCD Kcals consumed during early development (from birth through 16 months) predict brain structural alterations in adults. The sole use of females was another limitation of this study. Some human studies highlight sex differences whereby obese women, but not men, show structural brain alterations (Kroll et al., 2020). Future studies should include both males and females to assess whether social subordination stress- and diet-induced effects are different in males vs. females. Another limitation of this study is the lack of physiological or behavioral measures to understand the functional effects of the brain structural differences.

Our study not only sheds light on the unique effects of postnatal exposure to low social status and obesogenic diets on the structural development of neocortical and corticolimbic regions, but also on their persistence into adulthood. We identified developmental volumetric alterations in ICV and specific cortical regions due to the obesogenic diet consumption and/or social subordination stress. Notably, we identified unique life-long volumetric increases in cortical regions of SUB animals vital to social processing and cognition, that get magnified in late adolescence and early adulthood. Furthermore, these findings emphasize the possibility that these long-term differences in brain structure of individuals experiencing cumulative psychosocial stress (i.e. SUB) are neurostructural adaptations of the developing brain to increase survival in highly complex and challenging social environments. Additionally, our study highlights the brain’s reconstructive strength, as we found no significant long-term effects of obesogenic diets on structural brain development in adults. Ultimately, our findings emphasize the significant sensitivity and vulnerability of the developing primate brain and its reconstructive potential to social status and early postnatal consumption of obesogenic diets.

## ACKNOWLEDGEMENTS

This study was conducted with invaluable help from Natalie Brutto, Casie Lyon and additional research, animal care, colony management and veterinary staff at the Emory National Primate Research Center (ENPRC) Field Station, as well as from Ruth Connelly and Sudeep Patel at the ENPRC Imaging Center. This project was funded by NIH grants AG070704, HD077623, HD079969, MH078105-01S1, MH091645, U54 HD079124, the NIH’s Office of the Director, Office of Research Infrastructure Programs P51OD011132 (ENPRC Base Grant), and the EPC Fund for Excellence. The ENPRC is fully accredited by AAALAC, International.

